# Effect of Single Nucleotide Polymorphisms on the structure of long noncoding RNAs and their interaction with RNA Binding Proteins

**DOI:** 10.1101/2022.07.26.501647

**Authors:** Mandakini Singh, Santosh Kumar

## Abstract

Long non-coding RNAs (lncRNA) are emerging as a new class of regulatory RNAs with remarkable potential to be utilized as therapeutic targets against many human diseases. Several genome-wide association studies (GWAS) have catalogued Single Nucleotide Polymorphisms (SNPs) present in the noncoding regions of the genome, transcribing lncRNAs. In this study, we have selected 67 lncRNAs with GWAS-tagged SNPs and have also investigated their role in affecting the local secondary structures. The majority of the SNPs lead to changes in the secondary structure of lncRNA to a different extent by altering the base pairing patterns. These structural changes in lncRNA are also manifested in form of alteration in the binding site for RNA binding proteins (RBPs) along with affecting their binding efficacies. Ultimately, these structural modifications may influence the transcriptional and post-transcriptional pathways of these RNAs, leading to the causation of diseases. Hence, it is important to understand the possible underlying mechanism of RBPs in association with GWAS-tagged SNPs in human diseases.

## 1. Introduction

Nearly 90% of all the single nucleotide polymorphisms (SNPs), identified by Genome-wide association studies (GWAS), fall within the non-coding region of the genome, where apparently many lncRNA are synthesized (Giral et al., 2018; Mirza et al., 2014). LncRNAs are at least 200 nucleotides in length, seldom with an open reading frame of less than 100 amino acid length. Apparently, they are similar to mRNA in the context of the presence of 5’ cap structures and 3’ poly(A) tails (Zhang et al., 2014). They play a fundamental role in regulating various molecular pathways in several organisms. Their significance had been underestimated for a long time due to their non-protein-coding feature. However, recent research on lncRNA has completely reshaped their significance based on their function and localization. The functional attributes of lncRNA encompass its role as signal lncRNA (i.e., the transcription of lncRNA gives signaling cues for cellular processes), decoy lncRNA (i.e., titrates away the RBPs, transcription factors, microRNAs etc from stimulating the effector), guide lncRNA (i.e., directs the transcription factors or proteins and localizes them at particular gene loci for carrying out specific function) and scaffold lncRNA (i.e., acts as central support for various protein complexes to proceed with its function of gene expression) (Wang and Chang, 2011). Considering the genomic location, lncRNAs are distinguished as sense (i.e., transcribed from the sense strand of a gene), antisense (i.e., transcribed from the antisense strand of a gene), intronic (i.e., transcribed from introns of protein-coding genes), overlapping (i.e., when the coding regions within an intron are contained in the lncRNA on the sense or antisense strand) and intergenic (i.e., transcribed from intergenic regions of a gene) lncRNA (Ma et al., 2015).

SNP-induced structural changes in the noncoding region of the genome are difficult to capture, as it does not code and affect the protein sequence. The changes or alterations, in the regulatory region of RNA, affect the entire ensemble of RNA structures culminating in several human diseases (Halvorsen et al., 2010a). GWAS data has located genetic risk variants in various neurodegenerative diseases like Alzheimer’s disease, Parkinson’s disease and multiple sclerosis (Beecham et al., 2013; Chang et al., 2017; Jiang et al., 2017; Jun et al., 2017; Li et al., 2016; Liu et al., 2018, 2017d, 2017c, 2017b, 2017a, 2016, 2012; Zhang et al., 2015, 2018). Most SNPs associated with genetic elements are the cause of cancer, which is following the epidemiological observations of most familiar cancer risks (Gao and Wei, 2017).

The secondary structure prediction of the lncRNA sequence of interest indicated that some SNPs cause large-scale variation in their secondary structure. These structurally disrupting SNPs are called as RiboSNitches (Halvorsen et al., 2010b). RNA Binding Proteins (RBPs) govern the functional aspects of RNA through its specific binding sites (Guttman and Rinn, 2012; Wang and Chang, 2011). Recent evidence supports the fact that the interaction between lncRNA and RBPs regulates transcriptional events (Guttman and Rinn, 2012; König et al., 2011; Ulitsky and Bartel, 2013; Wang and Chang, 2011). HOTAIR lncRNA stands as an example to drive this point as this lncRNA helps in the recruitment of polycomb repressive complex 2 (PRC2) to specific mammalian loci to repress gene expression and promote cancer metastasis (Rinn et al., 2007). Recent investigations have also explored the functional annotation of lncRNA and RBP binding interaction but only to a limited extent since it is a herculean challenge to study the interaction of innumerable lncRNA and RBPs.

Some lncRNA employ regulatory complexes through its association with RNA binding proteins or RBPs and henceforth govern the expression of neighbouring genes (Engreitz et al., 2016). Thus, any kind of structural alteration to SNP containing lncRNA is likely to cause an impact on the molecular pathways and gene expression pattern in an organism leading to various disease conditions. Deciphering the mechanism, related to SNP containing lncRNA in regulating various molecular pathways through its interaction with RBPs, remains a prime challenge.

In this study, we have investigated the structural impact of SNP on lncRNA using an in-silico approach. Moreover, the effect of SNP on the thermal stability of the lncRNAs was measured. Further, we have analyzed how these structure disrupting SNPs alter the interaction pattern of lncRNA with RBPs at the level of sequence as well as structure level.

## 2. Materials and Methods

### 2.1 Mining of SNP containing lncRNA from lncRNASNP2 database

A comprehensive collection of SNPs containing lncRNAs are present in the lncRNASNP2 database (http://bioinfo.life.hust.edu.cn/lncRNASNP/#!/). The database provides extensive information on lncRNA containing SNPs and the impact of these SNPs on the structure and function of respective lncRNA (Miao et al., 2018). A file containing a total of 2109 GWAS tagged SNP was downloaded from the database. A comprehensive screening of the 2109 GWAS tagged SNPs was fetched us with 152 SNPs with P values less than 0.2 and located in the GWAS linkage disequilibrium (LD) region. Here P-value is a value of significance that distinguishes SNPs with major local effects on the functional and structural aspects of lncRNA and is computed by the RNAsnp program (Sabarinathan et al., 2013). The SNPs, found in the LD region, are those SNPs that have alleles correlated to alleles of neighbouring SNPs in the same chromosome of the genome. These SNPs were named tagged SNPs as they tag the location or capture the variation of nearby SNPs of a genome in governing a particular trait (Li et al., 2008). These tagged SNPs were further examined to find whether these SNPs containing regions are present within the local dissimilarity interval of lncRNA. The local interval is the region with maximum dissimilarity between the wild-type and mutant lncRNA. Careful scrutiny led to the filtration of 83 such structurally disrupting SNPs with local dissimilarity effect. SNPs present in the same/similar local interval sequence with different lncRNA ID were filtered out to finally give us 67 SNPs used in this study.

### 2.2 Obtaining minimal sequences of SNPs containing lncRNAs

The minimal sequence of SNP containing lncRNA or the sequence confined to the local interval region of lncRNA in the lncRNASNP2 database were obtained from the Ensembl Genome Browser (http://www.ensembl.org/) (Birney et al., 2004). In certain cases, the database links more than one lncRNA ID with the same SNP ID based on the consensus position of the interval sequence and their location in the same chromosome. An understanding of such puzzling cases of selecting a sequence was solved by analyzing the lncRNA ID in the NONCODE database (http://www.noncode.org/introduce.php) (Bu et al., 2012). Subsequently, distinct features such as exon number were examined for the lncRNAs from which it originates. Aspects like whether the lncRNA arises from the positive or negative strand of the concerned gene were also considered. The sequence was accessed by putting information about the local interval sequence in the search text box along with the selection of +1 or -1 strand in a particular format, i.e., Chromosome 10: 6,351,288 -6,351,335 as an example. Then this sequence was scanned against a complete sequence of lncRNA in NONCODE and lncRNASNP2 linked database (dbSNP). The consensus sequence present in both the databases was considered for further analysis. The sequence was manually mutated based on the allele position of SNP, as is presented in the lncRNASNP2 database. Hence the wild-type and mutant sequence of lncRNA was designed.

### 2.3 Structure prediction of lncRNA through various structure prediction software

LncRNA sequence was used to predict the structure and to understand the functional dimension of the sequence of interest. Although there is various secondary structure prediction software available online, the emphasis was given mainly to RNA structure (http://rna.urmc.rochester.edu/RNAstructureWeb/), RNAFold (http://rna.tbi.univie.ac.at/) and CentroidFold (http://www.ncrna.org/centroidfold/), as the parameters involved in different programs of concerned interest in these servers are novel with better accuracy (Bellaousov et al., 2013; Gruber et al., 2008; Sato et al., 2009). MaxExpect program of RNA structure was used to determine the secondary structure of lncRNA (Lu et al., 2009) employing default parameters. This program maximizes the accuracy of expected base pairs formed based on partition function, calculated on base-pair probability. One of the vital programs of the Vienna RNA package is RNAfold. It computes the minimum free energy (MFE) of the secondary structure of RNA (Zuker and Stiegler, 1981) and also gives base-pairing probabilities based on the partition function algorithm (McCaskill, 1990). Centroid structure was considered in our study for further analysis and evaluation. The framework of CentroidFold in predicting the secondary structure of RNA via RNA sequence input was established by applying the stochastic suboptimal folding algorithm like Sfold (Ding et al., 2005) and stochastic traceback algorithm for the CONTRAfold model as an alternative to the maximum expected accuracy (MEA) estimator of CONTRAfold (Do et al., 2006).

### 2.4 Identification of Protein binding sites on lncRNA

The database of RNA-binding protein specificities (RBPDB) (http://rbpdb.ccbr.utoronto.ca/) and RBPmap (http://rbpmap.technion.ac.il/), a computational tool, were used to identify the protein binding sites on lncRNA. RBPDB is a literature-based repository of experimentally identified RBPs in 4 metazoan species i.e., human, mouse, fly and worm. A total of 1487 *in vitro* and *in vivo* experiments on 1171 proteins are included in the database as per the latest release. A total number of 73 binding profiles in the pattern of position weight matrices (PWMs) are also included in the database. This certainly helps the user in determining the potential binding site of RBPs in the scanned lncRNA sequence by applying PWMs that compute the score of every single nucleotide at its position using Bioperl (Stajich et al., 2002) and generates a relative score that reflects the percentage of the score for a maximum score of the PWM computed. A threshold cut-off of the relative score for each site in the lncRNA sequence, which is greater than 80%, is reported and thus allows the user to identify the potential RBPs that bind to lncRNA prominently (Cook et al., 2011).

RBPmap predicts the binding site of RBPs on any lncRNA sequence by allowing the users for selecting motifs from a collection of 94 human/mouse and 51 *D. melanogaster* RBPs, which are experimental based and drawn out from literature as either a consensus motif or a Position Specific Scoring Matrix (PSSM). Accuracy and reliability of RBPmap were established by scanning the query sequence against a collection of RBPs in the database while computing a match score for the motifs in the lncRNA sequence and quantifying it in terms of a Z score coupled to P-value, calculated on Weighted Rank score (WR score) (Paz et al., 2014). The minimal sequence of lncRNA was scanned in both the databases and a list of human proteins was found that was showing binding specificity to the lncRNA sequence of interest. The heterogeneity was filtered out and a list of consensuses RBP, which showed binding specificity with the lncRNA sequence in both the database, were assembled. A sum of 26 such RBPs was enlisted in the databases, indicating their binding specificity with the lncRNA sequence.

### 2.5 Generating lncRNA and Protein 3D structure models

RNAcomposer (http://rnacomposer.cs.put.poznan.pl/) was used to generate a 3-dimensional (3D) structure of RNA from RNA secondary structure information (Biesiada et al., 2016). The interactive mode of RNA in the software was employed by entering the lncRNA sequence of interest and lncRNA secondary structure information in the form of dot-bracket notation. lncRNA secondary structure information in the form of dot-bracket was based on the centroid structure of the RNAfold web server. The Protein Data Bank (PDB) structure of lncRNA, generated in RNAcomposer, was downloaded and visualized in Pymol for clarity. The 3D structure of most of the RNA binding proteins was available in PDB Database (https://www.rcsb.org/) and their PDB structures were downloaded (Burley et al., 2019). RBPs like ELAV like RNA binding protein 2 (ELAVL2), serine and arginine splicing factor 13A (SFRS13A), and serine and arginine splicing factor 9 (SFRS9) were considered for homology modelling using Modeller 9.22 (https://salilab.org/modeller/9.22/release.html) (Eswar et al., 2007), as the experimentally determined structures were not available in PDB Database.

### 2.6 Docking of lncRNA structure against RBPs

Overall, 67 modelled structures of lncRNA were docked against specific RBPs to find out the difference between preferences of RBP for Wild type (WT) and SNP harbouring lncRNA on a structural context on the MPRDock webserver (http://huanglab.phys.hust.edu.cn/mprdock/). MPRDock server is based on the concept of efficient docking of RNA against protein ensembled by considering the flexibility of protein structures generated through homology modelling (He et al., 2019). Each docking run sample is in N × 4392 docking modes, where N is the number of proteins in the ensemble and 4392 corresponds to rotation of the sample in Euler space with an angle interval of 15 degrees. This sampling of binding modes in 3D Rotational space was preceded by a translation that corresponds to the best shape complementarity score. This was followed by scoring the binding modes based on Binding energy scores and ligand Root Mean Square Deviation (RMSD) values. A model with a better ligand RMSD value, out of the top 10 models of Protein-RNA complex in the output, was selected for our study.

## 3. Results

### 3.1 SNP-mediated structural variations in lncRNA

The lncRNASNP2 database was examined to investigate various aspects of GWAS-tagged SNP harbouring lncRNA. Table 1 depicts the location of a set of 67 structurally disrupting SNPs. The minimal sequences of all 67 lncRNAs were found in Ensembl Genome Browser and these sequences were manually mutated based on the location of SNP to design Wild-type (WT) and mutant (MUT) sequences or SNP harbouring sequences. RNA Secondary structure prediction servers, like RNAfold, RNA structure and CentroidFold, were employed to investigate the effect of SNP on the structural dimension of lncRNA. The results of these servers give us an insight into the role of some SNPs in disrupting the structure of lncRNA. It is observed that although the algorithms and parameters used in these 3 specific servers for structure determination were essentially different to a certain extent, yet calculation of lncRNA folds based on partition function in these three servers, produces consensus results for the lncRNA. Figure 1A elucidates the schema of lncRNA secondary structure with paired and unpaired regions highlighted in blue and red colours respectively. Figure 1B represents the percentage change in the structured region from WT to MUT lncRNA with emphasis on the differences in the output shown as the error bars calculated from the three servers in use. The quantitative degree of change in the structured region was calculated to be zero in a few lncRNAs in the presence of SNP. A prominent feature of these SNPs, except for rs6952808 and rs569214, was the type of mutation, i.e., transition mutation; where a pyrimidine ring was replaced with another pyrimidine ring, or a purine ring was replaced with another purine ring. These substitutions are less likely to alter the structure of lncRNA (Lyons and Lauring, 2017). Similarly, the quantitative difference between the structure of WT and MUT lncRNA reflected as consensus prediction from the employed servers was moderate to major in SNPs rs10743430, rs115199861, rs115199861, rs76037120, rs7837688 respectively as shown in table 1.

**Table 1.**
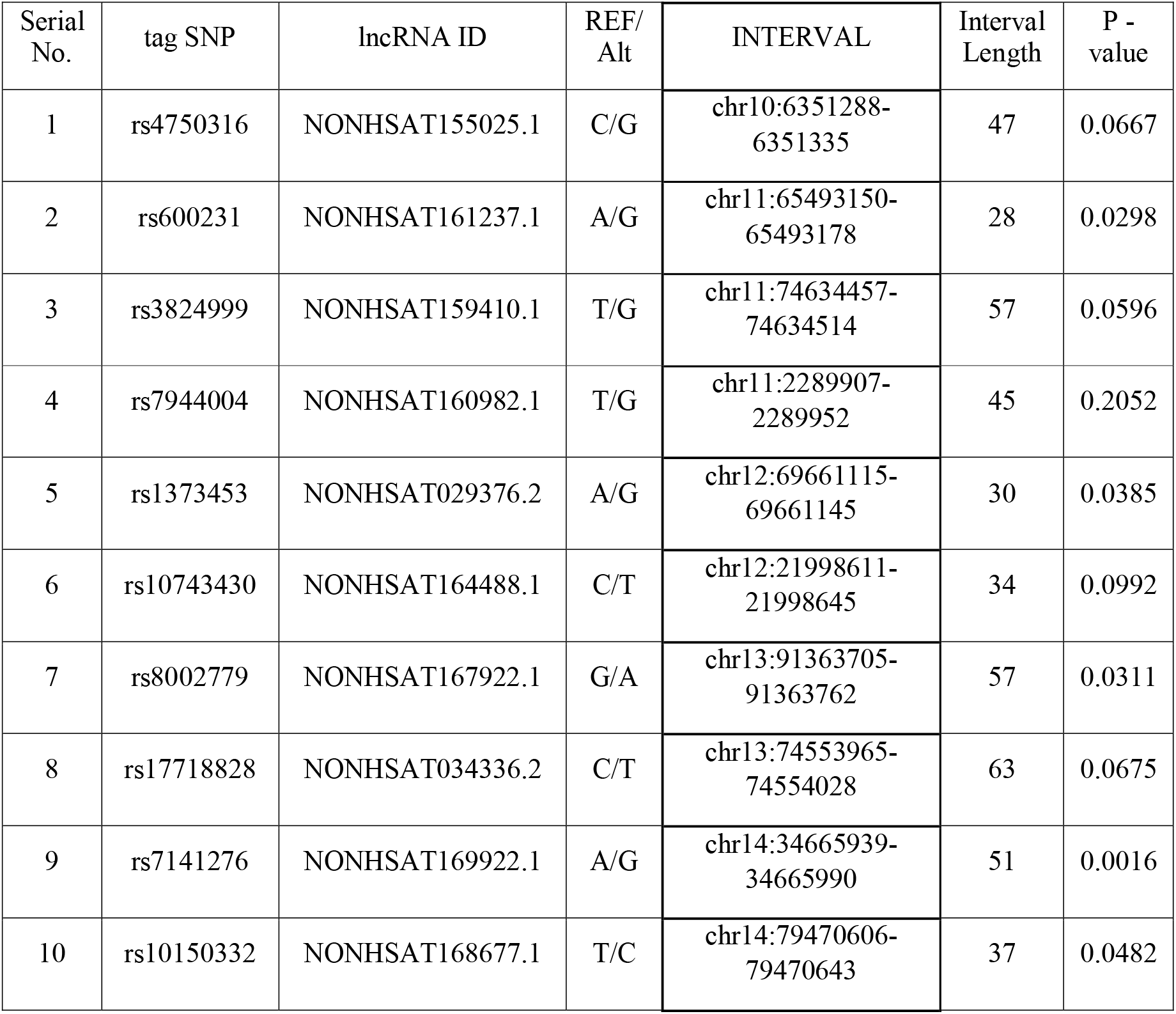
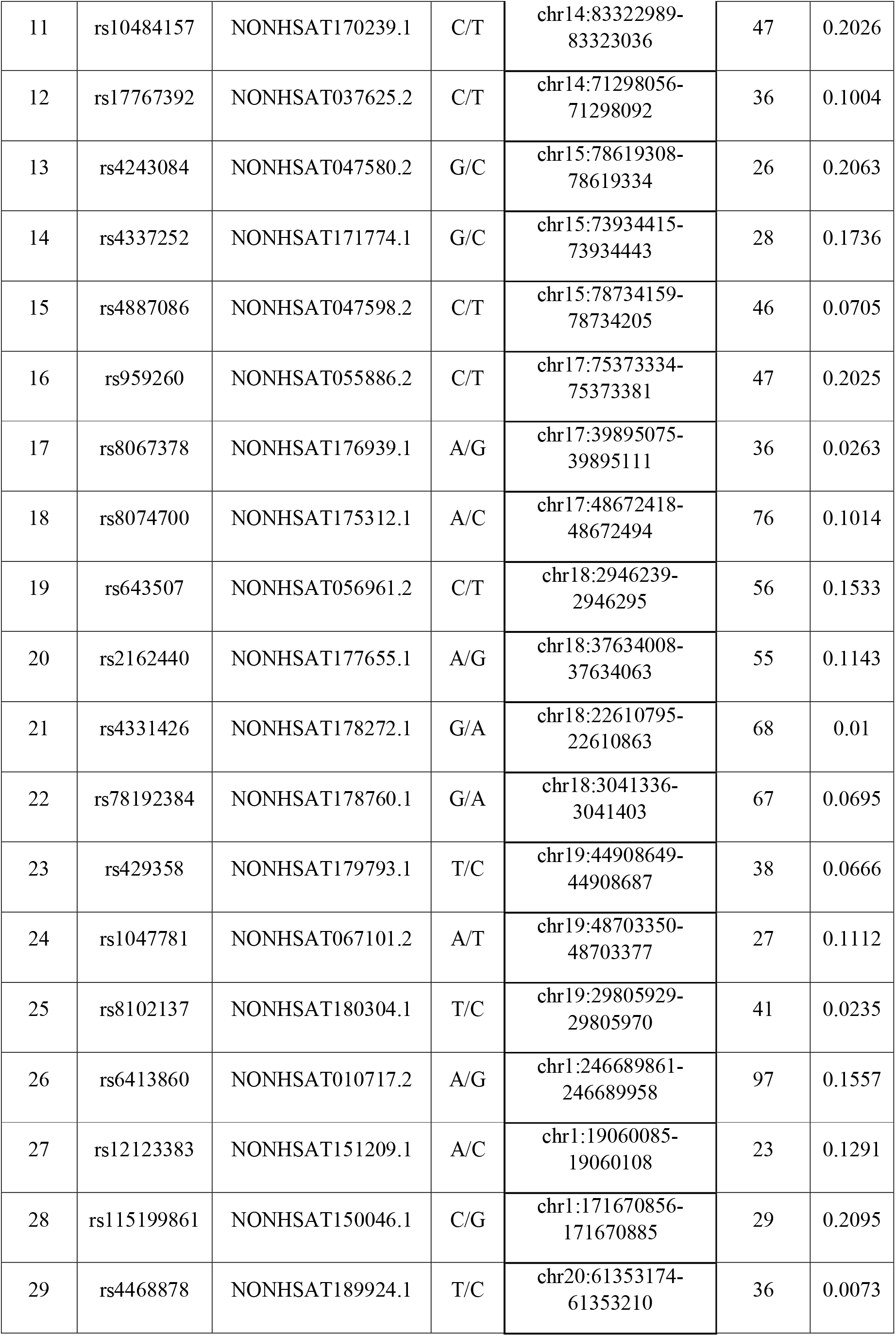
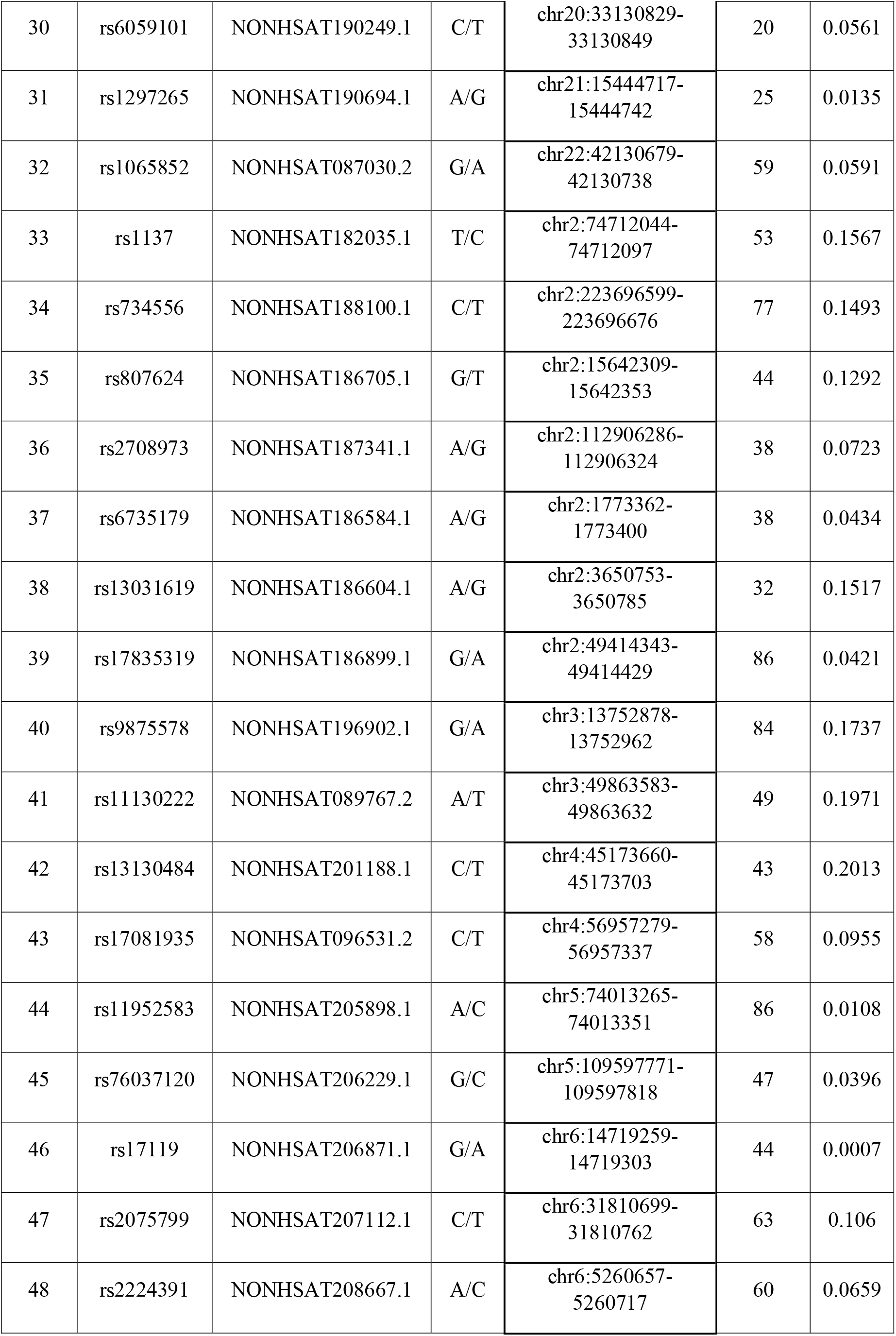
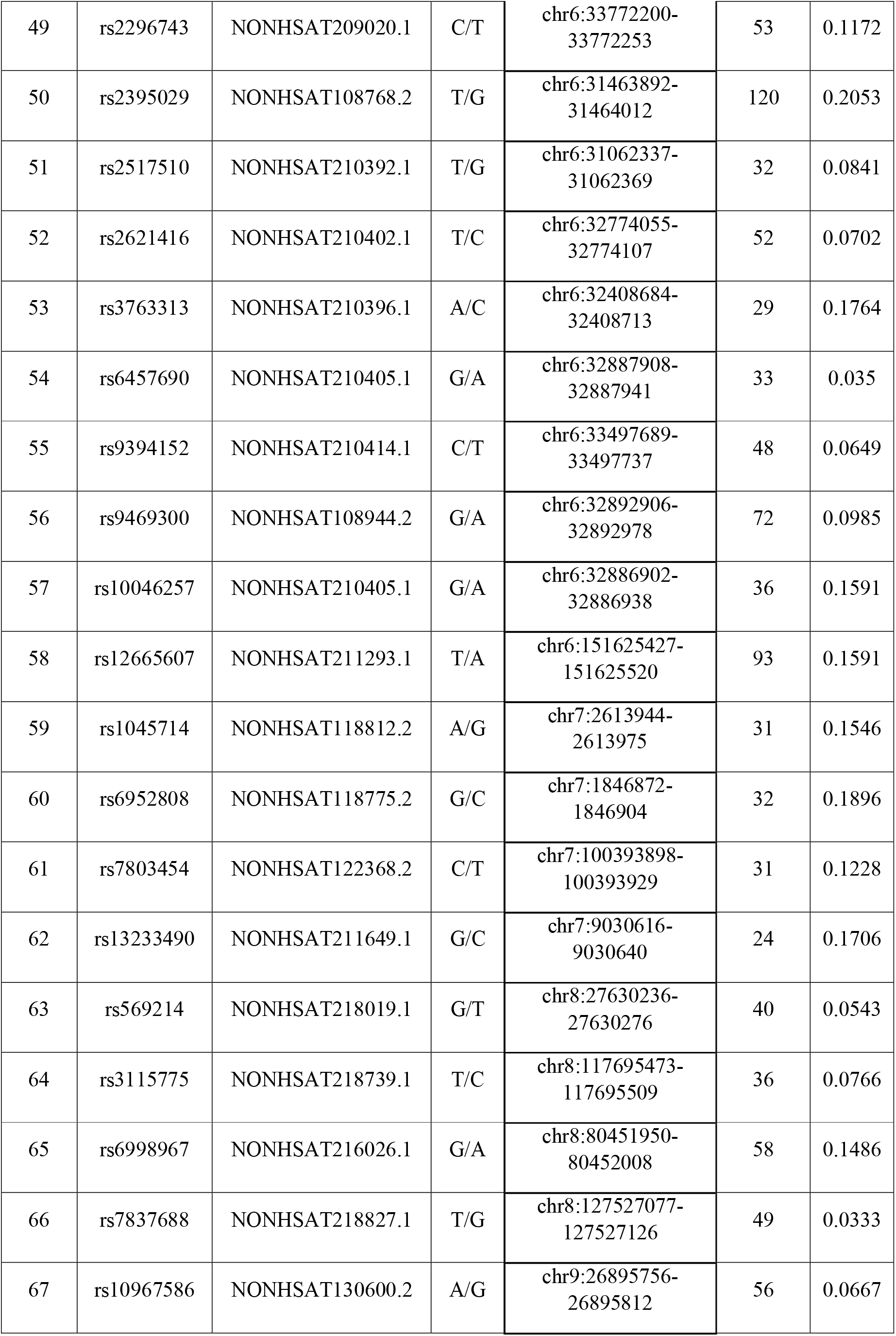
SNP ID of tagged SNPs along with lncRNA ID where these SNPs are presented Interval length in the sequence length with maximum dissimilarity region used for the analysis.

**Figure. 1.**
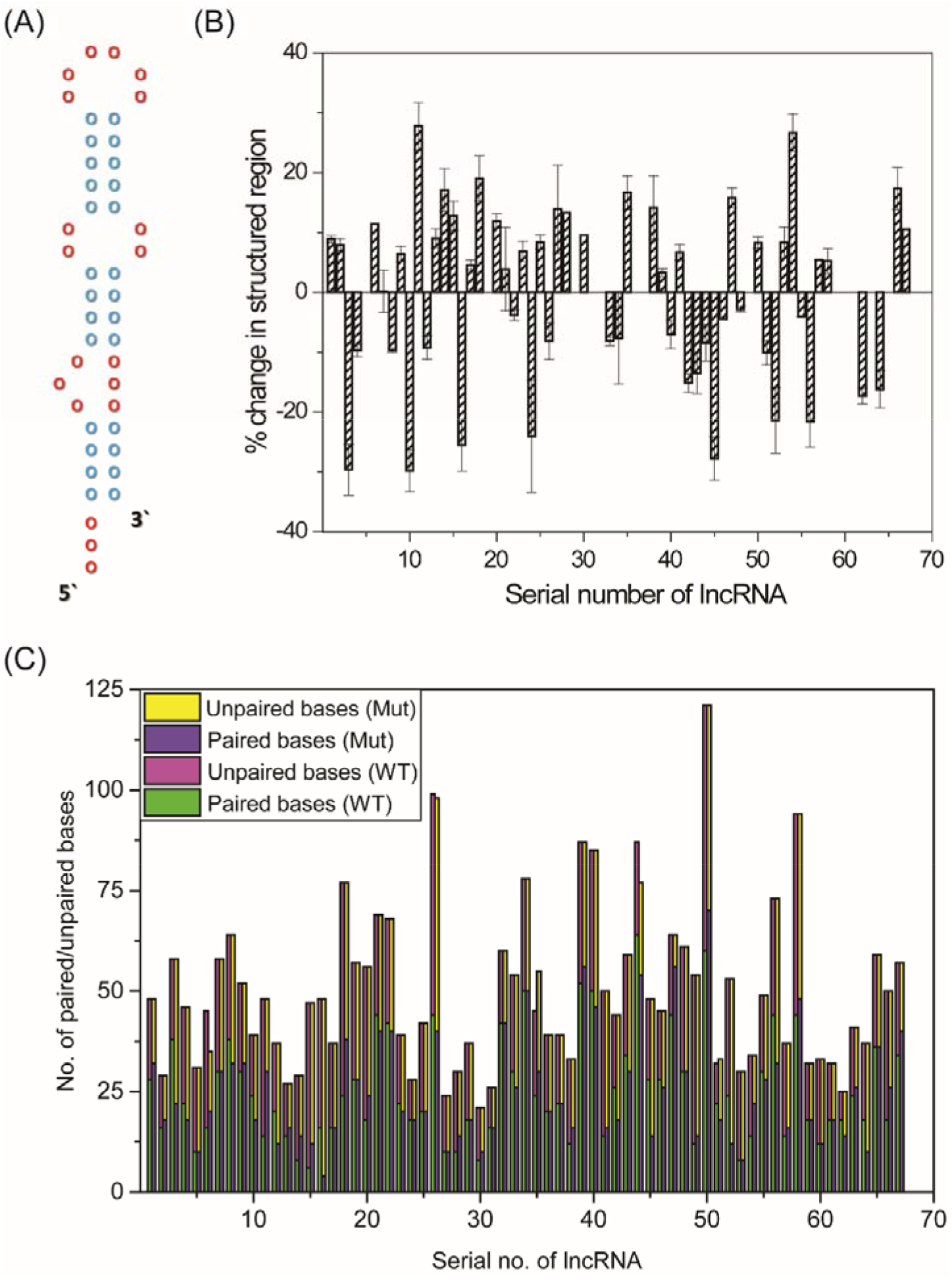
Structural analysis using three web servers for predicting the secondary structure of lncRNA. (A) The schema of lncRNA structure with paired and unpaired regions (red circles represent unpaired bases and blue circles represent paired bases). (B) The percentage of secondary structural change in lncRNA as calculated by the number of base-paired/unpaired changes in the structure of WT to MUT lncRNA. The error bars represent the standard deviation of the output of 3 employed structure prediction servers in specifying the percentage change. (C) The marked difference in base pair probability matrix based on Euclidean distance for all 67 lncRNAs.

### 3.2 Nucleotide Sequence determining the propensity of RBP towards lncRNA

The mapping of the binding propensity of RBP with the lncRNA sequence of interest was evaluated by RBPDB and RBPmap. The consensus RBP hits, found in both databases, were considered for further analysis. The RBP interacting, with the lncRNA sequence of interest in RBPDB, had a relative score greater than 80%. Similarly; the RBP found, as hits in RBPmap for a particular lncRNA sequence of concern, were evaluated based on a Z score coupled to a P-value against an algorithm of the weighted rank score (Paz et al., 2014). Table 2 illustrates a set of 24 different RBPs that have a binding affinity with 67 different lncRNA with due consideration to WT and SNP harbouring or MUT sequence.

**Table 2.**
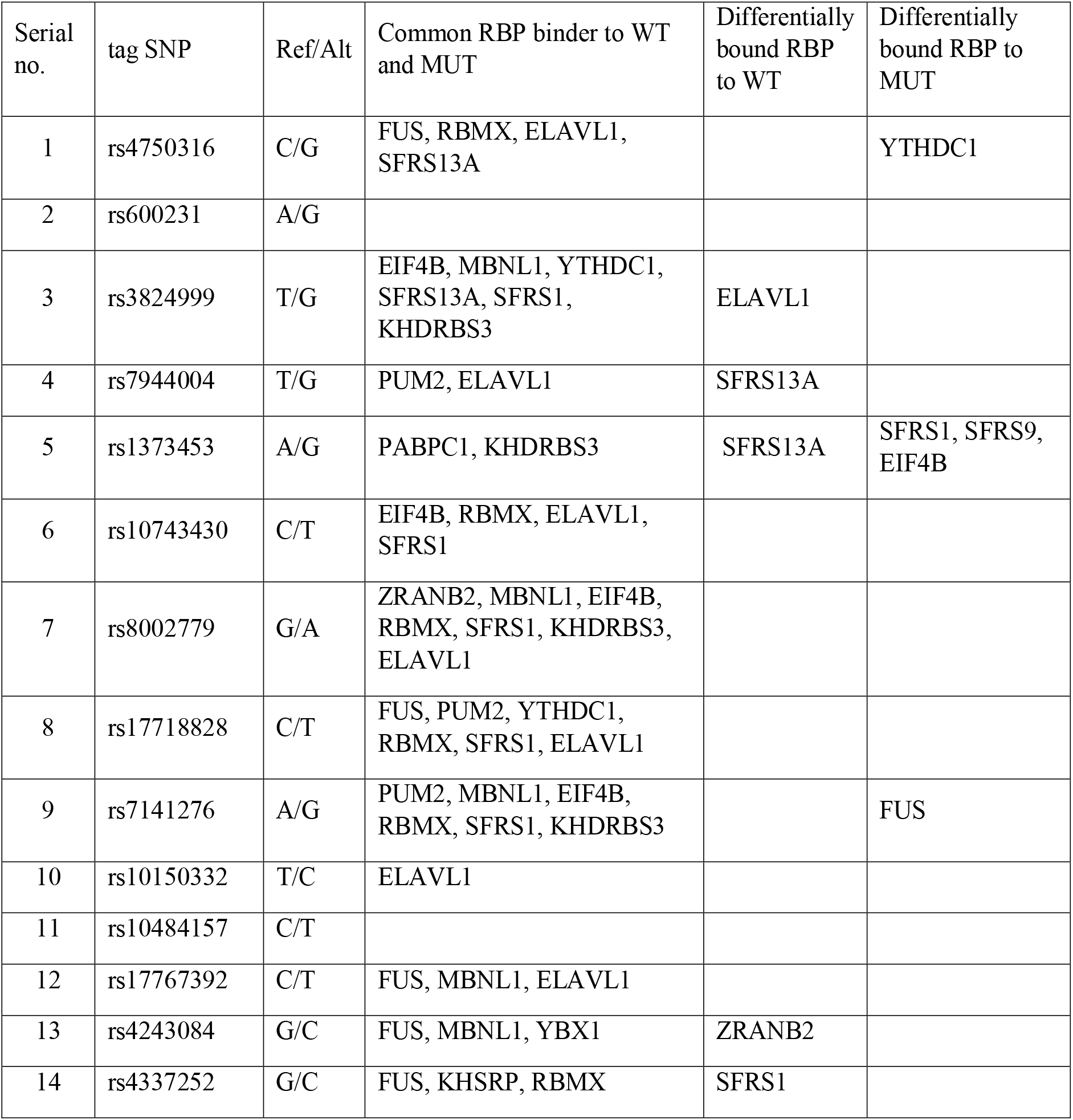
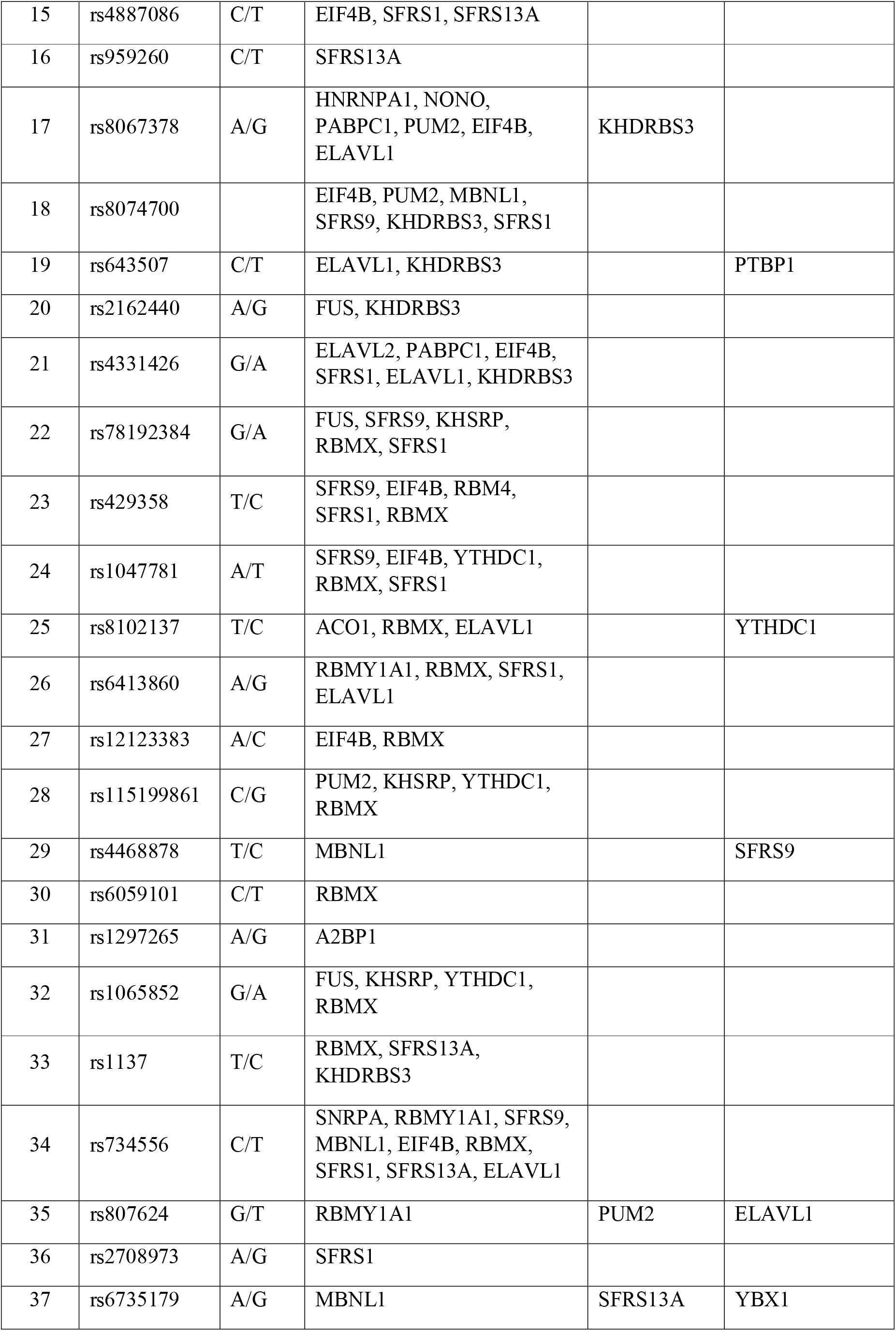
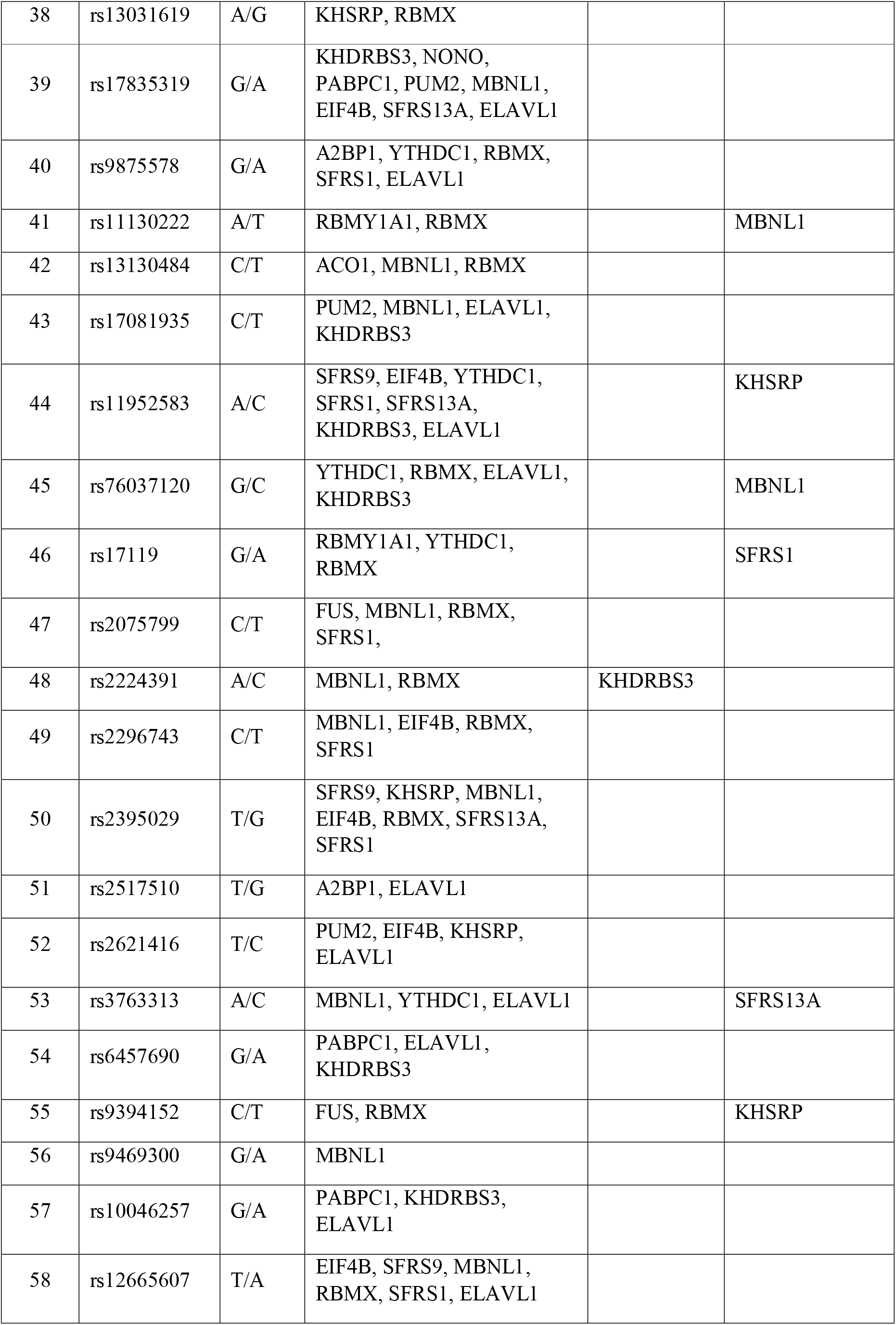
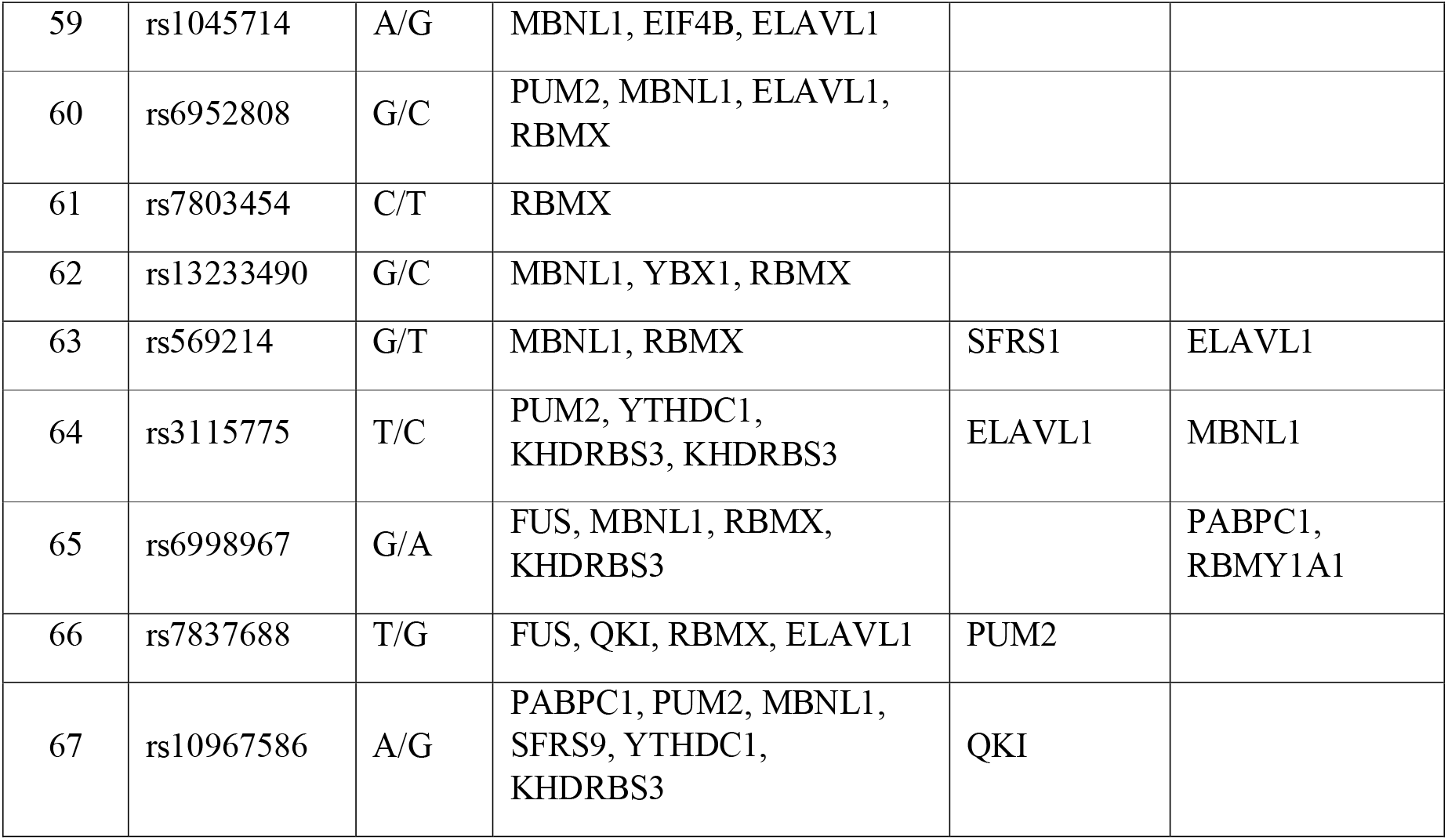
Consensus RBPs binding to WT and the SNP sites in lncRNA was obtained from RBPDB and RBPmap.

Figure 2 represents a schema of how different RBPs interact with lncRNA sequences based on a particular motif. Mostly RBPs with RNA recognition motif (RRM) and K Homology (KH) domains, were found as hits in both the databases when scanned against a lncRNA sequence of interest. General lncRNA binding RBPs: like RNA binding motif protein, X Chromosome (RBMX), Muscleblind like splicing regulator1 (MBNL1), Fused in sarcoma (FUS), KH domain-containing, RNA-binding, signal transduction-associated protein 3 (KHDRBS3), ELAV like RNA binding protein 1 (ELAV1) and serine and arginine splicing factor 1 (SFRS1) were also found as hits in both databases. These RBPs are splicing regulatory proteins that commonly regulate the function of most lncRNAs (Gerstberger et al., 2014). There were also cases of specific RBPs like Quaking Homolog, KH Domain RNA Binding (QKI), Non-POU Domain Containing Octamer Binding (NONO), Aconitase1 (ACO1), ELAV-like RNA binding protein 2 (ELAV2), RNA Binding motif Protein 4 (RBM4) and Ataxin 2-binding protein 1(A2BP1) that bind to very few lncRNA in a sequential context. This might be due to their specific functions. For example, ACO1 is important for maintaining the level of iron in cells (Kaptain et al., 1991); 8041788, (Stehling et al., 2013). Similarly, QKI and A2BP1 are involved in maintaining neuronal plasticity (Åberg et al., 2006; Gaudet et al., 2011). An anomaly was observed for the lncRNA sequence with SNP ID rs600231 and rs10484157, for which none of the RBP was found to have the binding propensity as per the RBPDB database. Further, several RBPs were found as hits for their binding propensity to these two lncRNA sequences of interest in RBPmap but with optimal Z score (−1 to +1) coupled to suboptimal P values (>0.005) or suboptimal Z score (−3 to +3) coupled to optimal P values (<0.005). Hence such RBPs were not considered for further evaluation. It was observed that in some cases SNP alters the binding propensity of RBP to a particular lncRNA sequence. Concentrating on consensus results produced from structure prediction servers, it was found that transition mutation that substituted A with G in the MUT sequence, changed its binding motif for a maximum number of RBPs for lncRNA with SNP ID rs1373453 (Figure 2A). Hence, it was established that for this lncRNA, SFRS13A was differentially bound to WT lncRNA sequence and SFRS1, SFRS9 and Eukaryotic translation initiation factor 4B (EIF4B) were differentially bound to MUT lncRNA sequence, since the binding motif <u>A</u>GAC in WT lncRNA favoured its affinity for SFRS13A and the altered motif <u>G</u>GAC in MUT lncRNA favoured its affinity for SFRS1, SFRS9 and EIF4B. Similarly, Polypyrimidine tract binding protein 1(PTBP1) was found to be differentially bound to MUT lncRNA with SNP ID-rs643507 due to altered motif <u>U</u>CU instead of <u>C</u>CU in WT lncRNA sequence (Figure 2B) and SFRS9 was found to be differentially bound to MUT lncRNA with SNP ID rs4468878 due to altered motif AGCA<u>C</u> instead of AGCA<u>U</u> in WT lncRNA sequence (Figure 2C).

**Figure. 2.**
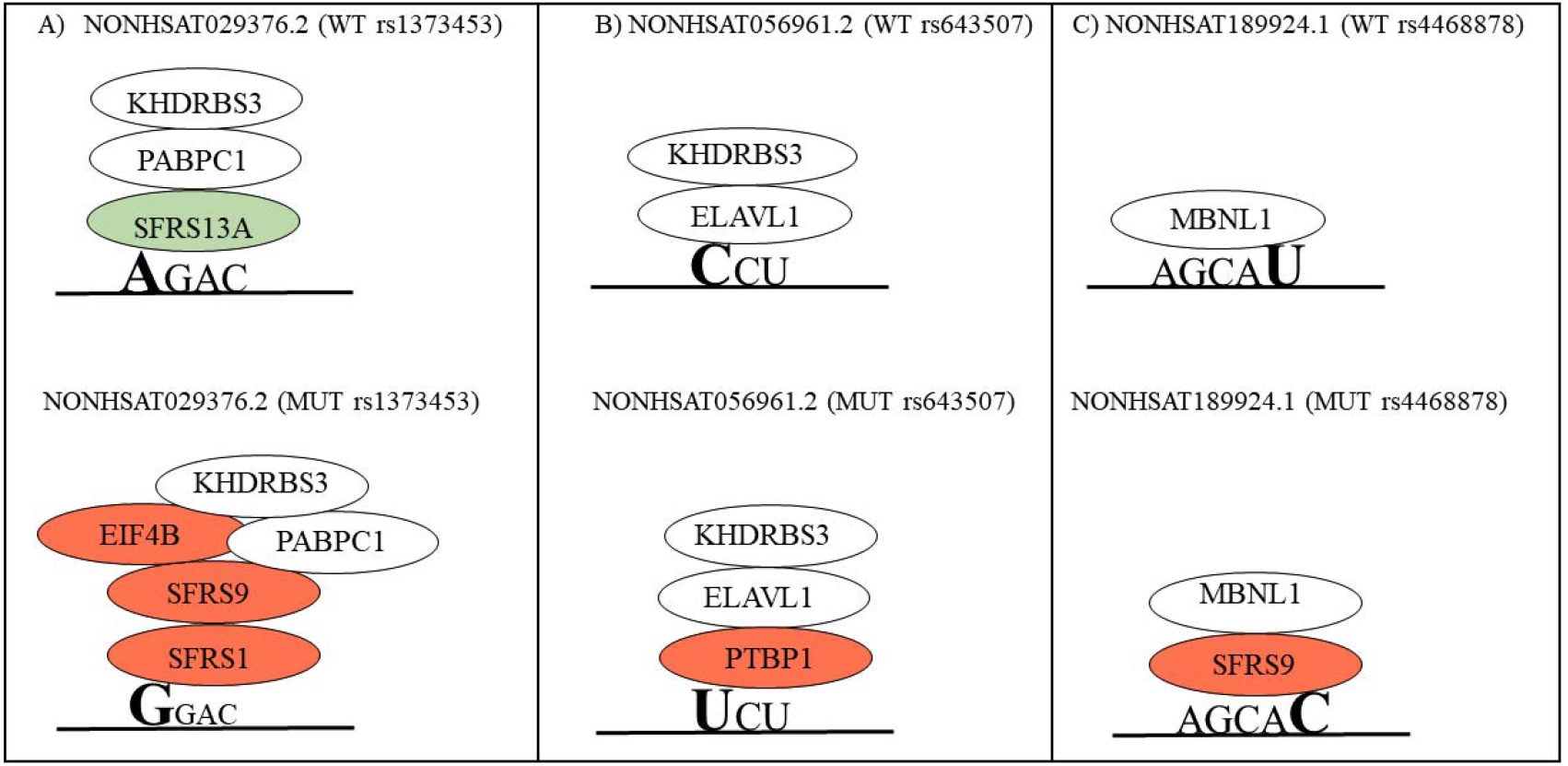
Analysis of RBPs binding to lncRNA sequence and representative schema for different RBPs recognizing the SNP region (WT and MUT) in lncRNA sequence motif of (A) rs1373453 (B) rs643507 (C) rs4468878.

### 3.3 lncRNA structure plays an important role in regulating the binding propensity of RBPs

It is speculated that SNP harbouring region in a lncRNA sequence may not fall in the RBP binding region in most cases, yet SNP might alter the structure of lncRNA or disrupt the RBP binding motif. Figure 3A elaborates the Delta Docking scores and highlights significant differences in the binding affinity of some RBP when docked against WT and MUT lncRNA. This difference in docking score might be due to profound secondary structural alteration in WT lncRNA in the presence of such SNP. In certain cases, it was discovered that RBP like QKI, which plays an important role in neuron plasticity and development, binds differentially to WT and SNP harbouring lncRNA with SNP ID rs10967586 in a sequential context. However, when QKI was docked against the same lncRNA, no such differential docking score was found as per the results (Δ docking score = 17.5), which showed nearly the same Docking score and ligand RMSD values. RBMX binds most of the lncRNA in a sequential context and is a splicing regulatory protein. However, as per our docking results, RBMX was found to bind differentially to WT and SNP harbouring lncRNA in some cases where there was a huge difference in structure between WT and SNP harbouring lncRNA. Figure 3A elaborates the docking scores that range from -200 to -300 in the majority of the lncRNA-RBP complexes and highlights significant differences in binding affinity of some RBP when docked against WT and MUT lncRNA. RBP like RBMX and A2BP1 showed a higher binding affinity with MUT lncRNA of SNP ID rs9875578 when the delta docking score was significantly large with values -598 and -545 respectively. Similarly, another significant delta docking score was seen for lncRNA with SNP ID rs11130222 when the WT and MUT lncRNA were docked against RBMY1A1 and RBMX protein with the Δ docking score of -607.7 and -790.9 respectively. This might be due to profound structural alteration in WT lncRNA in the presence of SNP. Figure 3B illustrates that the delta docking score was regulated by different domains of RBPs when docked against WT and MUT lncRNA. The results also illuminate us with the knowledge of various domains of RBP in regulating the function of lncRNA and it was noticed that the RRM domain has a substantially higher binding affinity to lncRNA. Figure 3C sheds light on RBPs that may be considered as general RNA binders since they bind with most lncRNA irrespective of the presence of SNP. The pie chart in Figure 3C represents RBPs binding to the selected 67 lncRNAs and gives an idea about the selective or general binding nature of RBPs. Some of the RBPs like RBMX, ELAV, MBNL, and KH-Type Splicing Regulatory Protein (KHSRP) appear to be binding to most of the lncRNAs suggesting their general binding nature and some of them including Y-box binding protein 1 (YBX1), QKI, A2BP1, NONO, ACO1 seem to be selective binders. The docking score also displays an important feature of RBPs, signifying that the RRM domain of RBPs has the capacity to buffer the effect of the changes made by SNPs (Maris et al., 2005). In many cases, the docking scores for WT and MUT lncRNA are nearly the same even in the presence of structurally disrupting SNPs or riboSNitches.

**Figure. 3.**
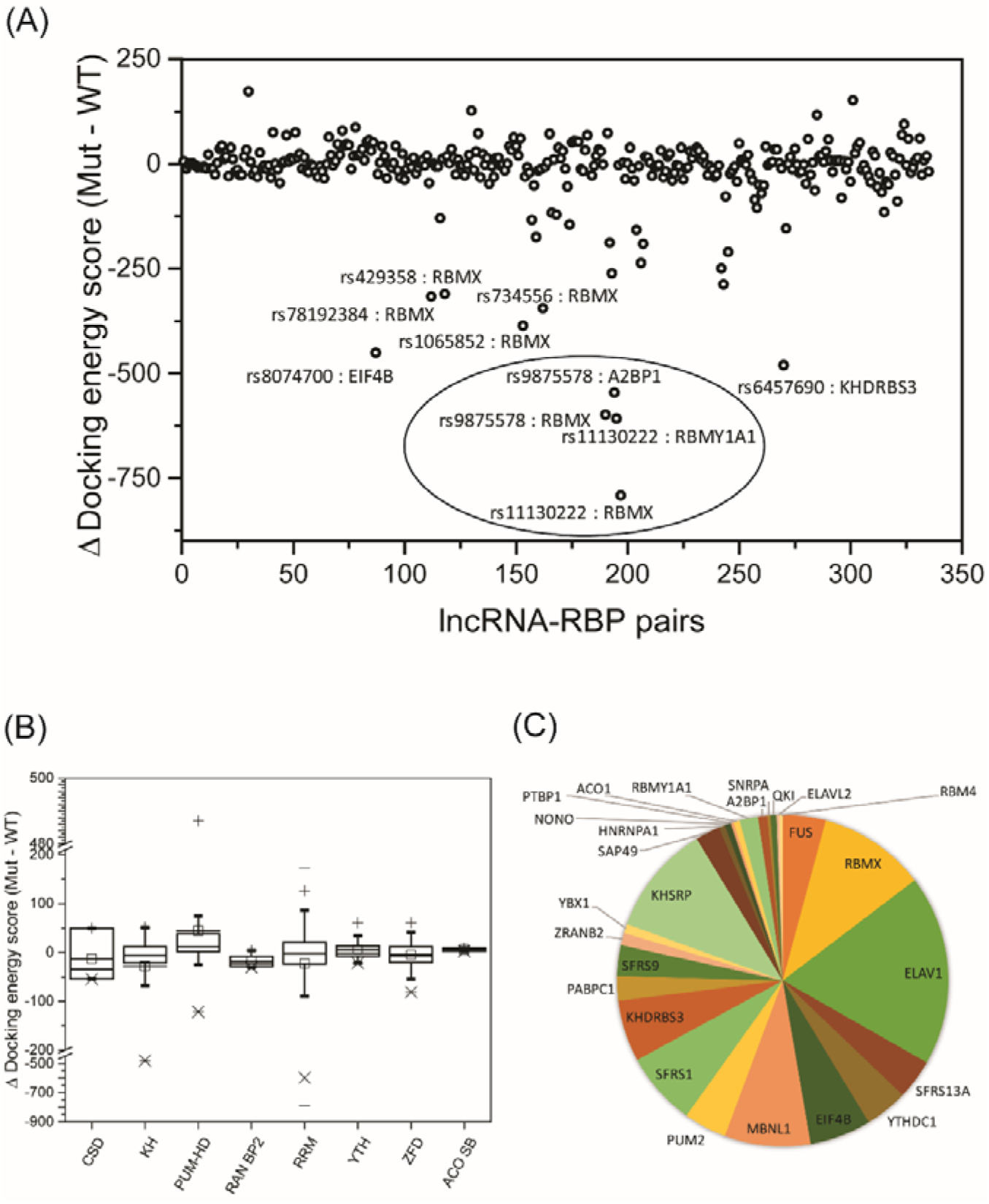
Molecular docking analysis of lncRNA WT and MUT binding with different RBPs. (A) Delta docking score for all 67 lncRNA bound to RBPs. (B) Regulation of Delta docking score for all 67 lncRNAs by different domains of enlisted RBPs. RRM domain of RBPs show higher affinity for RNA-RBP interaction. (C) Regulation of docking of the lncRNA-RBP complex by different RBPs on a quantitative basis. The RBPs bind to lncRNAs irrespective of their structural changes with the help of RRM domain.

### 3.4 Differential Docking score of RBMX and A2BP1 when docked against WT and MUT lncRNA with SNP ID rs9875578

A clear distinction was seen between the WT and SNP harbouring lncRNA with SNP ID rs9875578 in RNAFold. The arc plot representation of these two structures is visibly distinct as shown in Figure 4A. When the A chain of RBMX (Moursy et al., 2014) as receptor protein was docked against the structure of WT and SNP harbouring lncRNA of interest, the binding affinity score or the van der Waals interaction score for WT and SNP harbouring lncRNA was -409.29 and -1007.89 for A chain of RBMX. This shows that SNP alters the fold of specific lncRNA such that a general binding RBP like RBMX shows a strong affinity for MUT lncRNA as compared to WT lncRNA. The complex structure of WT and MUT lncRNA with RBMX protein is shown in Figure 4B. The ligand RMSD value, which shows the deviation of conformation for lncRNA structure in bound form from a native form, is less in the case of SNP harbouring lncRNA as compared to WT lncRNA.

**Figure. 4.**
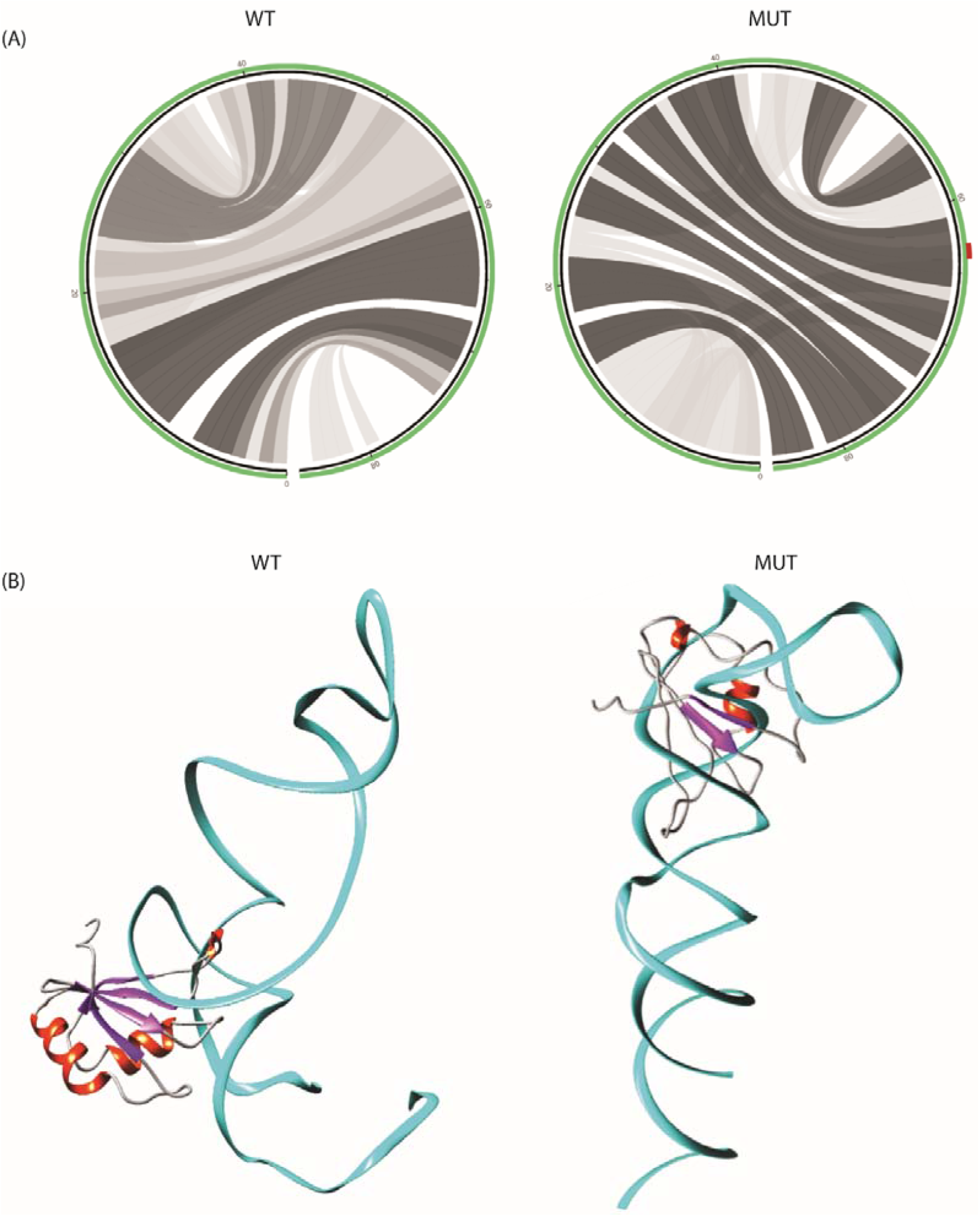
Structural change between WT and MUT lncRNA and their interaction with the same RBP. (A) Arc plot elucidates the structural change for lncRNA serial no.40. This structural dissimilarity is due to the presence of SNP in the MUT type. (B) It shows the complex structure of lncRNA no. 40 with RBMX. There is a difference in the lncRNA-RBP interaction, and this is because of low binding affinity score of MUT type. It represents that MUT type has more affinity for RBMX as compared to WT type.

## 4. Discussion

Attention and research on Protein–centric approach to determine the structural and functional attributes of the RNA-Protein complex has already reached a height of evolution. Turning our glance, through the unexplored domain of RNA-centric approach to meet the understanding of RNA-Protein complex, needs an intense amassing of data from in silico and in vitro techniques. The purpose of our investigation to break the enigma surrounding, the lncRNA-Protein complex via an exploration of lncRNA sequence and structure on a preliminary in silico scale, has fetched us certain perplexing yet promising results. The effect of Single Nucleotide Polymorphism (SNP) on mRNA has been studied to a certain extent and the consequences of such SNP on RNA-Protein complex have been dealt with curiosity and credibility (Shen et al., 1999). In comparison to the mRNA sequence, the lncRNA sequence is less conserved. However, the structure of lncRNA, shown to be conserved, plays a vital role to provide a mechanistic explanation of the functional dimension of SNP-affected lncRNA (Derrien et al., 2012).

The computational RNA structure prediction server points out that there is no such universal approach to deduce the structure of lncRNA as its structure is flexible with multiple degrees of freedom across a plane (Duarte and Pyle, 1998; Gendron et al., 2001; Hershkovitz et al., 2006; Leontis and Westhof, 2001; Murray et al., 2003). Yet certain homogenous output across all the servers directed our attention to some lncRNAs with tagged SNPs. Reverting to previous research done on SNPs, it was expected that transversion is more structurally disrupting as compared to transition, since they affect the regulatory functions of RNA profoundly, even forming the ground for non-synonymous mutation of amino acids (Guo et al., 2017). Although structural disruption for SNP harbouring lncRNA (rs6952808) is due to Transversion (G28C), an unexpected, calculated output of minor entropy change of 0.079 and distance of 0.0001 between WT and MUT base-pairing probability, from Mutational Analysis of RNA (MutaRNA) server, displays important characteristics of transversion. Transversion shows a major structural disrupting effect based on nearest neighbour nucleotides present adjacent to SNP in +1 as well as in -1 direction and if it affects the helix of RNA or the single-stranded loops and bulges (Zhao and Boerwinkle, 2002). In the studied case, SNP causes an alteration in the motif UGC and forms a new motif UCC. The nucleotide bases, present adjacent to the SNP, are pyrimidines and not a heterogeneous combination of A and T, with the single-stranded location which might not show a pronounced effect of structural disruption. Hence, the resultant output shows a new direction around the spectrum of Transversion which might not show intense structural disruption owing to certain conditions. Most of the tagged SNPs are case of transition that causes synonymous or silent mutation of amino acids. A confounded attribute of transition was also considered in this study and most of the lncRNA, undergoing a change in the base pairing network, were contributed to transition, as predicted in various RNA structure prediction servers.

This estimated negative correlation between lncRNA sequence and structure was established on the grounds of SNP, which revealed the phenomenon of metabolic processes in our body that could be significantly affected by major structural disruption in lncRNA. In this study, the noncoding RNA was explored, which may not have an impact on the formation of protein. But it might have a huge impact on its affinity and specificity for RBPs owing to its regulatory role played to influence epigenetic changes in cells. Hence, the resulting outcome of the servers has enlightened us with the fact that all SNPs may not be structurally disrupting to a significant extent as they might lead to a labile and unbalanced cellular process in our body progressing into various diseases (Glinsky, 2008).

The relevance of results, found in RBPDB and RBPmap, gives due consideration to the SNP harbouring lncRNA sequence and showcases the idea that SNP falling outside the binding motif of RBP does not impact its affinity for the altered lncRNA sequence. But SNP falling within the binding motif of RBP alters its affinity for the lncRNA sequence. If this was the case, there would be large disturbances in the pathways or mechanisms in our body regulated by the lncRNA–RBP complex. Hence, here comes the role of lncRNA structure as a deciding factor in guiding the formation of the lncRNA-RBP complex. If we give equal weightage to the docking results, then the structural aspect of lncRNA, in forming a strong complex with its partner RBP, cannot be ignored. Commonly RBPs, that regulate splicing, bind to lncRNAs irrespective of the presence of SNPs. This might be due to the common buffering mechanism shared by these common RBPs. Even in some cases, a cooperative effect could be seen for these RBPs while binding to a particular lncRNA. This cooperation is facilitated by lncRNA structure that allows an assembly of proteins on lncRNA in a synergistic or mutually exclusive event (Bartel, 2004; Komili et al., 2007; Somarowthu et al., 2015) and posttranscriptional gene regulation from genome-wide studies to principles (Landthaler et al., 2008). Although some RBPs are sequence-specific and most of them have the RRM domain, there might be a deviation from this assumption based on research. Some of these RBPs with RRM domains bind to various RNA through sequence specificity as well as structure affinity (Heym and Niessing, 2012; Müller et al., 2011). This is the case for most splicing regulatory RBPs like ELAV1, RBMX and MBNL1, that bind to these 67 lncRNAs irrespective of the presence or absence of RBP. So, the docking scores are nearly the same for most of the lncRNA bound to RBP. However, an SNP affecting lncRNA sequence might not alter its conformation tremendously to minimize the impact on its affinity for binding partner Protein.

Some proteins with unique and vital functions have a differential binding affinity with particular lncRNA based on structural disruption by SNP. It enhances the knowledge of the functional aspect of the particular RBP and lncRNA. lncRNA and the partner RBP, with correlated function, lead to this differential affinity of RBP with SNP harbouring lncRNA. From the result, it is found that lncRNA, harbouring SNP (SNP ID-rs9875578), binds strongly with RBMX as compared to WT lncRNA. It is due to the structural alteration, i.e., instead of ‘G’ nucleotide is replaced by ‘A’ in the MUT lncRNA. The WT lncRNA is expressed in the brain tissue (Cabili et al., 2011) and the SNP habouring lncRNA is associated with anxiety disorder (Coleman et al., 2016). The RBP RBMX is a genome stabilizing protein that prevents the process of cryptic splicing (Luzzi et al., 2020). It is associated with brain development in Zebrafish (Tsend-Ayush et al., 2005). Mutation of genes that depends upon RBMX for productive expression can cause mental retardation and other diseases like microcephaly (Genin et al., 2012; Pulvers et al., 2010). So, it may be suggested that WT lncRNA and RBMX play a vital role in the process of brain development while the interaction between MUT lncRNA and RBMX leads to brain-related anomalies.

## 5. Conclusion

The outcome of the in-silico work, when corroborated by in vitro and in vivo work, could give us promising output in identifying and understanding the various functions played by the lncRNA-RBP complex in our body and how the structure of lncRNA contributes significantly to its function. Further, it would also enhance our understanding of the role of the epigenetic mechanism behind various diseases; facilitating our path in a new direction of devising therapeutic targets for molecules to treat some of the dreadful diseases and possibly provide a plausible solution to reduce the impact of these diseases.

## Acknowledgments

Mandakini Singh acknowledges a junior research fellowship (F.No.16-9(June 2019)/NET/CSIR) from the university grant commission, India. S.K. acknowledges the research funding from SERB (EEQ/2018/000022), India.

